# Comparison of the human’s and camel’s erythrocyte deformability by optical tweezers and Raman spectroscopy

**DOI:** 10.1101/2022.08.02.502368

**Authors:** Tuna Pesen, Mete Haydaroglu, Simal Capar, Mehmet Burcin Unlu, Ugur Parlatan

## Abstract

The evolution of red blood cells (RBCs) or erythrocytes has led to variation in morphological and mechanical properties of these cells among many species today. Camelids have the most different RBC characteristics among the vertebrates. As a result of adaptation to the desert environment, camelid RBCs can expand twice as much of their total volume in the case of rapid hydration yet are almost undeformable under mechanical stress. In this work, the difference between cell features of the human and the camelid species was explored both mechanically and chemically with optical tweezers and Raman spectroscopy, respectively. We measured the deformability of camel RBCs relative to the human RBCs at the single-cell level using optical tweezers. We found that the deformability index (DI) of the camel and the human RBCs were 0.024±0.0188 and 0.215±0.061, respectively. Raman spectral analysis of the whole blood of these two species indicated that some of the Raman peaks observed on the camel’s blood spectrum were absent on the human blood’s spectrum, which further points to the difference in chemical contents of these two species.

## Introduction

All the vertebrates, except Antarctic icefish, have circulating hemoglobin packed in red blood cells. These cells are responsible for oxygen transport for all modern vertebrates.^1^ Among the vertebrates, RBCs vary in shape and size, ranging from over 50 *μ*m to below 10 *μ*m.^2^ The possible outcomes of such variations to RBCs’ mechanical functions are still under investigation.^3^ Regarding such variations in RBCs’ shape, size, and deformability, there is no unique description of optimized RBC characteristics.

Human RBCs are biconcave-shaped viscoelastic cells. Having an average diameter of about 8*μ*m^4^, they can deform through the small capillaries (with an inner diameter less than 3*μm*) without losing cell functions. Thereby, deformability is a crucial property for them to circulate through the vascular system effectively.

On the other hand, camelid RBCs are rigid and almost undeformable under mechanical stress.^5,6^ Rather, their ellipsoidal shape makes them aligned with the stress field. Thereby, they can transverse through the circulatory system owing to their shape.^5^ Yet, they can expand twice as much of their initial cell volume,^7–10^, which advantage camelids survive in the desert environment in the case of a large amount of hydration.^11–13^ Such expandability and, at the same time, low deformability of these cells is related to the cell membrane phospholipid composition, which makes the membrane more fluid and the cells resistant to osmotic variations.^11,14,15^ On the other hand, in the case of dehydration (water loss up to 27%), the fluidity of the camel blood is not affected since the camel erythrocytes make up a small amount of the blood content.^16^

The structural differences between the camelid and human RBCs, in terms of osmotic stability, was formerly explained by the protein-lipid ratio and the protein organization of the cell membrane.^17–19^ A study reported that the protein-lipid ratio for human RBC ghosts (which are dead cells without cytoplasmic structures) was measured as 1.25, while this value for camel RBCs was 3.0.^17^ Another study showed that the protein-lipid ratio was 1.2 and 3.7 for human and camel erythrocyte membranes, respectively.^18^ The same study also indicated that the amount of membrane integral proteins was almost five times greater in camel cells.^18^ In another study, high resistance to osmotic lysis of the camel erythrocytes was explained by changes in major integral proteins and the distribution of the glycoproteins in the cell membrane.^20^

Camel RBCs were investigated with varying techniques and modalities such as Klett photometer,^21^ microcinematography,^9^ flame photometer,^22^ sonic irradiation,^17^ gel electrophoresis,^6,18,20^ viscometric techniques,^5^ scanning electron microscope,^6^ scanning and transmission electron microscopy,^19^ chromatography,^23^ blood count,^12^ proteomics,^15^ and rheometry.^24^ These studies focused on osmotic behaviors, susceptibility to direct hemolysin of cobra venom, protein composition, and organization of the membrane, the relationship between cell shape and membrane structure, homeostasis, and viscosity.

Optical trapping is a handy tool for life sciences, primarily cell mechanics investigations. It enables fixation-free immobilization and manipulation of the biological materials in a desired way. The physical principles behind this tool rely on the momentum conservation of light when interacts with matter. The change in light’s momentum, due to the interaction with the particle of interest, forms optical forces on the particle.^25–28^ Controlling this forces on the particle, it is possible to perform cell manipulation such as stretching. When a correct wavelength is used, it has become possible to investigate a cell in its living medium with little or no photodamage. By adding a second beam or splitting the original beam into two, a dual-beam optical tweezers system can be created where the cell mechanics is interrogated by stretching cells.^29–32^ Double-beam optical tweezers studies have two types of geometry: i) the beams counter-propagate^32–34^ ii) both beams are in the axial direction and catch the cell from two ends.^35–39^ Among these studies, deformation of red blood cells was investigated for healthy cells,^34,35^ type 2 diabetes mellitus and diabetic retinopathy,^33^ and radiation therapy applied cells.^36^

As the mechanical properties of the cell change, the chemical structure also changes. An efficient way to monitor this change is Raman spectroscopy. This technique examines the vibrational transition energies due to laser-molecule interaction. The frequencies of the photons scattered from the sample differ from the illumination frequency as much as the molecular vibration frequency^40^ Raman spectroscopy was utilized to understand the mechanical and chemical structure of the red blood cells previously for some of the species but has never been applied to camel blood yet.^41–45^

In this study, we aimed to approach the camel RBCs from the perspective of cell mechanics by measuring cell deformability. For the deformability measurements, we used optical tweezers to apply direct mechanical stress on an isolated RBC. By trapping the erythrocytes from the two ends and moving one of the traps with a certain speed, mechanical stress was created, and cell stretching was achieved. In addition, for chemical analysis, we utilized Raman spectroscopy of whole blood samples from the two species.

## Methods

### Ethical Statement

With the ethical permission (2022-05) of Bogazici University Science and Engineering Fields Human Research Ethics Committee (FMINAREK), one healthy woman and two healthy men volunteered for this study. Experiments with two healthy male single-humped dromedaries were conducted with the permission (2021-017) of Bogazici University Institutional Animal Experiments Local Ethics Committee (BUHADYEK).

### Blood Collection

For the study of human erythrocytes, blood samples were drawn using a lancet needle from the participants’ fingertips. A veterinarian has drawn blood samples for the camelid RBC experiments and the samples were transported inside EDTA tubes at 4^°^C. All the measurements were completed on the same day of blood collection.

### Sample Preparation

Whole blood of 0.1 *μ*L was mixed with 1 mL PBS (Phosphate-buffered saline) and 100 *μ*L BSA (Bovine serum albumin) solution in a tube. 70 *μ*L of the prepared blood sample was placed onto the microscope slide, and BSA-dried cover glass was placed on top. The edges of the cover glass were sealed using nail polish. The BSA-dried cover glass was used to prevent RBCs from adhering to the cover glass. The BSA-dried cover glass was prepared by placing 40 *μ*L BSA on a cover glass and kept in an incubator at 45^°^C for 10 minutes. The experiments were performed at room temperature.

### Experiment and Analysis

The experiments were performed using dual-beam Zeiss PALM Micro Tweezers, with a wavelength of 1064 nm, and a laser power of 60 mW (30 mW for each trap). The determined stretching parameters (velocity, direction of movement and the power of the traps, and experiment duration) were set on the user interface of the tweezers. Before starting the experiment, the optical traps (while off) were configured on the cells. For human RBCs, the traps were positioned on the two ends of the cell with a trap separation of 5 *μ*m. Whereas, for camel RBCs, the traps were positioned at the center of the cells on top of each other. After the positioning procedure, the traps were activated, and the experiment was started. During the stretching, one of the traps was moving with the defined velocity of 1 *μ*m/s for 10 s, while the other one was kept fixed in position. When the trap started to move, the RBC first stretches, and then after reaching the maximum stretched length, it escapes from the moving trap and begins to relax. The experimental setup for optical tweezers and Raman spectroscopy can be viewed in Figure 1.

**Figure 1.**
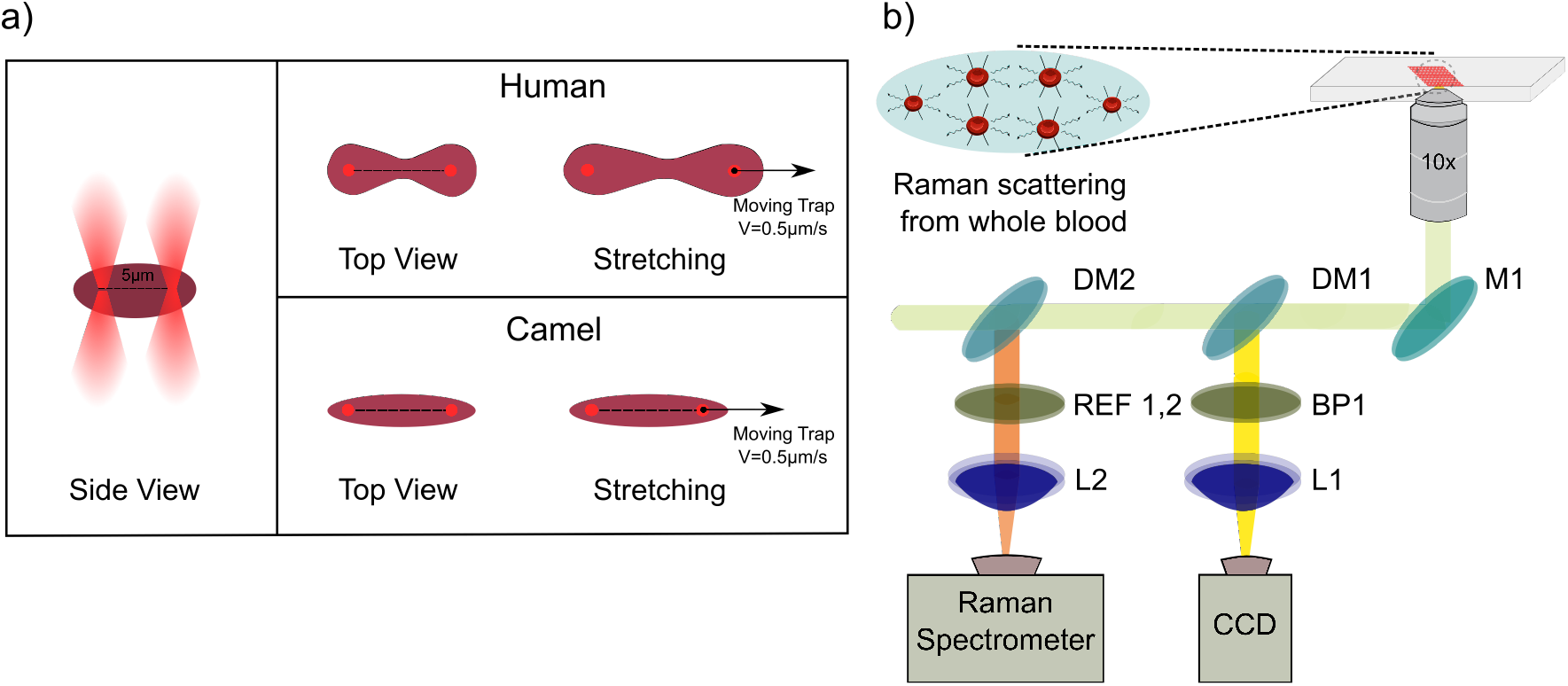
a) The cell stretching process at the single-cell level on optical tweezers setup, b) Demonstration of whole blood sample analysis on Raman spectroscopy setup.

Feret diameter is commonly used in method to analyze the size of cells.^46,47^ We used the maximum Feret diameter function of MATLAB^48^ to calculate the axial diameter of the erythrocytes. Deformability index (DI)^33^ was calculated using the following equation:

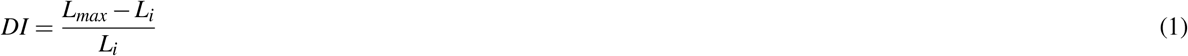

where, *L*_*i*_ is the initial (unstretched) length, *L*_*max*_ is the maximum stretched length of the RBC.

## Results and Discussion

In this study, we measured 206 human and 159 camel RBCs using optical tweezers. Single stretching-relaxation process of an individual RBC can be viewed in Figure 2.

**Figure 2.**
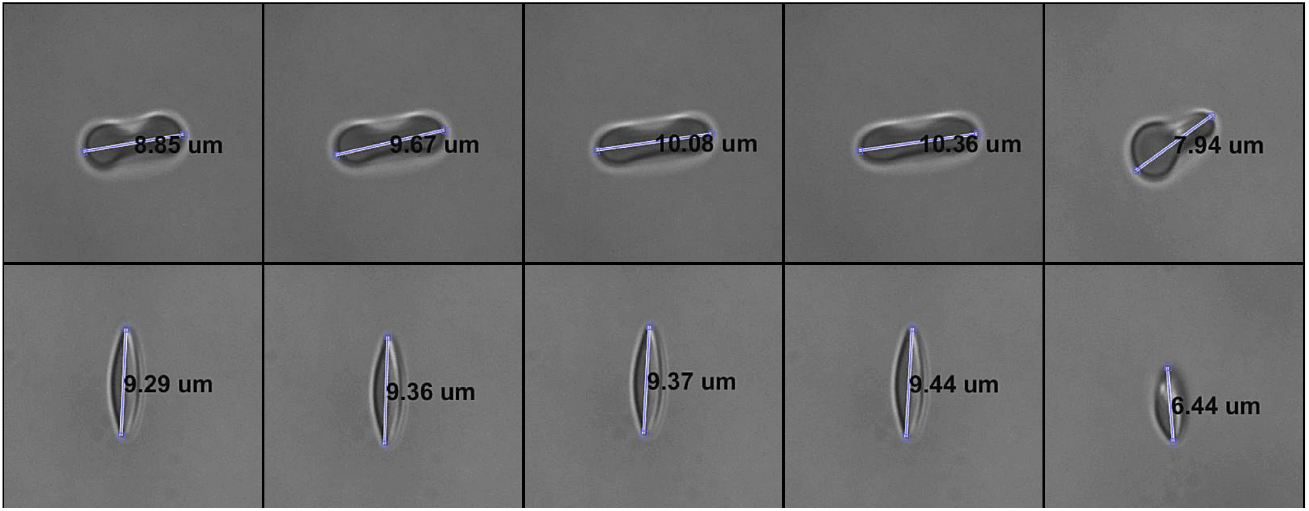
Top view of stretching and relaxation processes of a human (top row) and a camel (bottom row) RBCs. Calculated Feret diameters were labeled on the RBCs in each frame.

Firstly, we compared the initial, and the maximum stretched lengths of the cells. The kernel fit to the data revealed that kernel bandwidths for the camel group are 0.2964 and 0.3130, and for the human group, they are 0.2304 and 0.2496, respectively (Figures 3). While the separation of the L_*i*_ and L_*max*_ curves is very small in the camel group, these two curves have dramatically differed from each other in the human group. Such a result supports the current literature, which points out that the camel RBCs are less deformable than the human RBCs.

**Figure 3.**
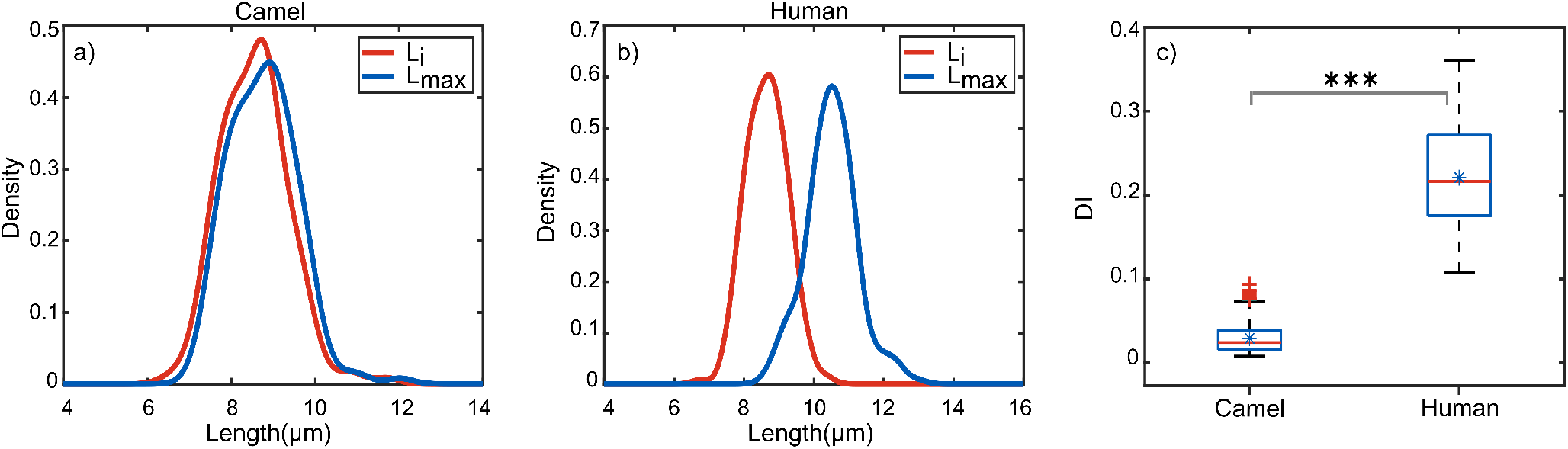
Normal kernel distribution of initial and maximum stretched lengths with the corresponding bandwidths: a) for the camel 0.2964 and 0.3130 and, b) for the human 0.2304 and 0.2496 c) Boxplot of the DI. Mean values and the standard deviations of DI for camel and human groups were calculated as 0.024±0.0188 and 0.215±0.061, respectively. The result of the t-test was p<0.001.

In Figure 3c, we demonstrated the boxplot of the deformability index of the RBCs calculated using the Equation 1. The graph revealed the mean DI of the camel and human groups with the standard deviations as 0.024±0.0188 and 0.215±0.061, respectively. This result showed that camel RBCs are almost one order of magnitude less deformable than human RBCs. The student’s t-test was applied to the DI data and the difference was found as statistically significant (p<0.005).

To obtain chemical information from RBCs of the two species, we acquired Raman spectra of the whole blood of the two groups. We repeated the measurements with the Raman spectrometer ten times for each blood sample to decrease error. In Figure 4 we demonstrated the difference between the average Raman spectra of both species. The average Raman spectra are different in several Raman shift positions. To quantify the difference and to reveal the sub-components of the important peaks, we applied a Gaussian curve-fitting on the spectra. The main differences between the two species are located in the Raman shift values stated in Table 1.

**Figure 4.**
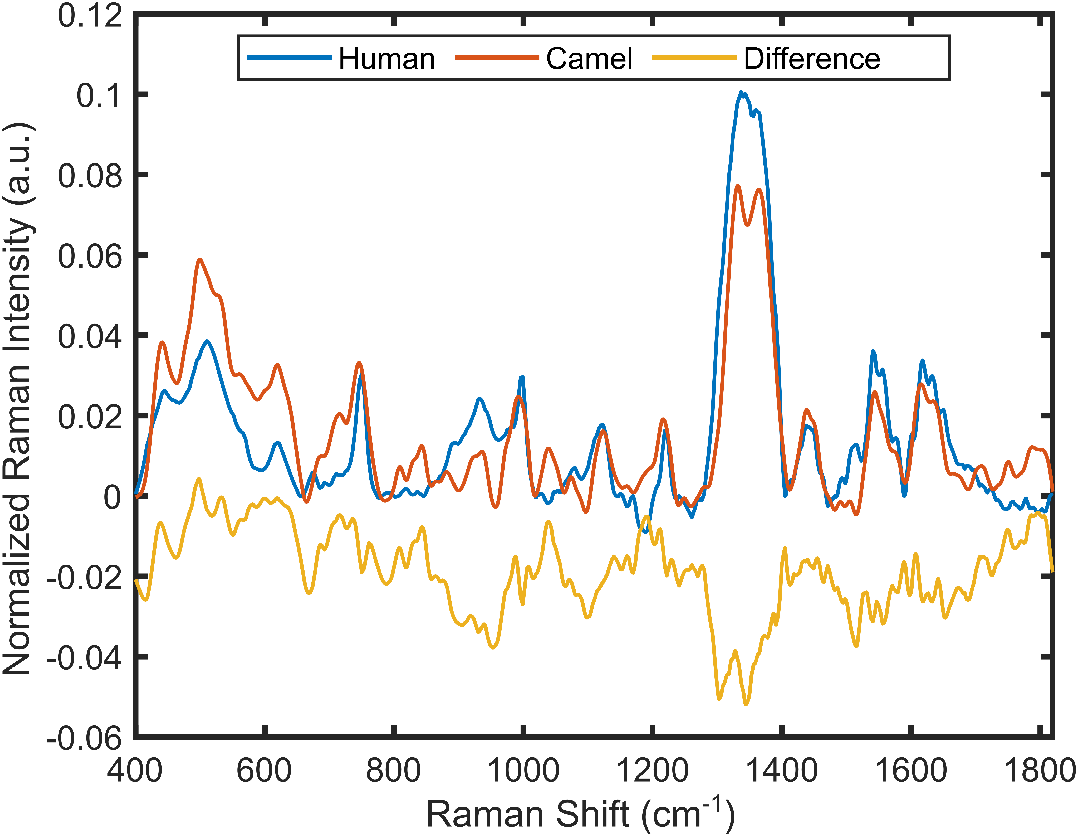
Raman spectrum of the two groups indicates dramatic differences in some bands’ intensities: 540 cm^−1^, 714 cm^−1^, 1080 cm^−1^. The bands, 838 cm^−1^, and 945 cm^−1^, are absent on the human spectrum, while these bands are Raman-active on the camel spectrum.

**Figure 5.**
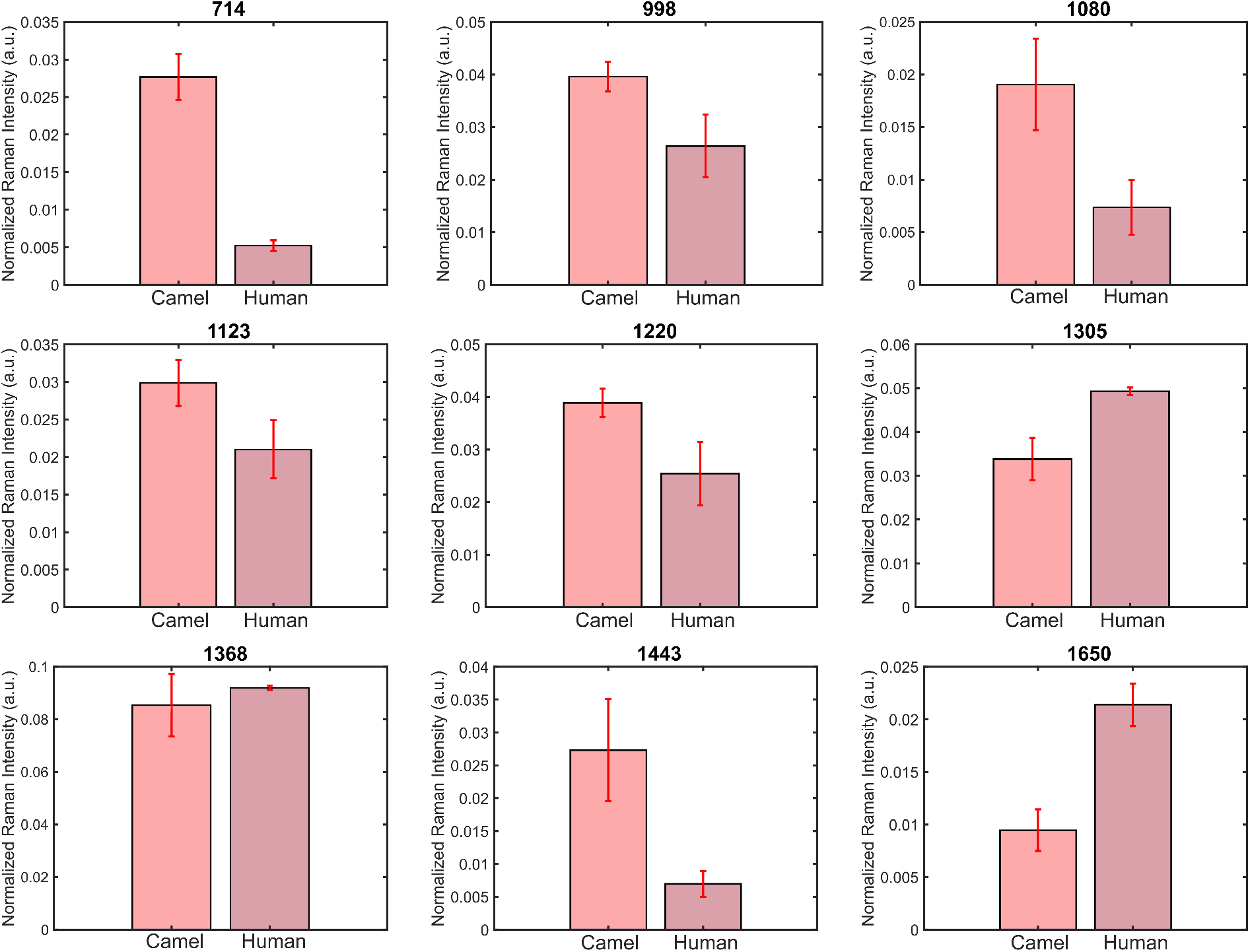
The bar graph shows the selected peak Raman intensities after band component analysis. These peak intensities are correlated to the amount of lipid and protein compositions; 714 cm^−1^ and 1080 cm^−1^ phospholipids, 1220 cm^−1^ Amide III, 1305 cm^−1^ and 1443 cm^−1^ lipids, and 1650 cm^−1^ Amide I.

**Table 1.** The band assignments of human and camel RBCs’ Raman spectra.

| Raman Shift($cm^{-1}$) | Normalized Raman Intensity (a.u.) | | Assignment |
| --- | --- | --- | --- |
|  | Human | Camel |  |
| 540 | $0.0022 \pm 0.0078$ | $0.0042 \pm 0.0009$ | S-S<sup>49,50</sup> |
| 714 | $0.0052 \pm 0.0007$ | $0.0277 \pm 0.0031$ | C-N (phospholipid membrane)<sup>51</sup> |
| 838 | absent | $0.0303 \pm 0.0017$ | Tyrosine<sup>52,53</sup>, Deformative vibrations of amine groups<sup>54</sup> |
| 945 | absent | $0.0144 \pm 0.0034$ | $\delta(\text{pyr})_{\text{asym}}$, Skeletal modes (polysaccharides, amylose)<sup>49,55</sup> |
| 998 | $0.0264 \pm 0.0026$ | $0.0396 \pm 0.0028$ | $\nu(C_{\beta}C_1)_{\text{asym}}$<sup>56</sup> |
| 1080 | $0.0074 \pm 0.0026$ | $0.0190 \pm 0.0043$ | $\nu(C_{\beta}C_1)_{\text{asym}}$, phospholipids<sup>56,57</sup> |
| 1123 | $0.0047 \pm 0.0025$ | $0.0300 \pm 0.0031$ | $\nu(\text{pyr half-ring})_{\text{asym}}$<sup>56</sup> |
| 1220 | $0.0254 \pm 0.0061$ | $0.0388 \pm 0.0027$ | $\delta(C_mH)$, Amide III<sup>56</sup> |
| 1305 | $0.0493 \pm 0.0009$ | $0.0338 \pm 0.0049$ | $\delta(C_mH)$<sup>56</sup> |
| 1368 | $0.0920 \pm 0.0008$ | $0.0854 \pm 0.0119$ | Heme, $\nu(\text{pyr half-ring})_{\text{sym}}$<sup>56</sup> |
| 1439 | $0.0161 \pm 0.0006$ | $0.0273 \pm 0.0078$ | $\delta(CH_2/CH_3)$<sup>58</sup> |
| 1557 | $0.0326 \pm 0.0058$ | $0.0343 \pm 0.0023$ | $\nu(C_{\beta}C_{\beta})_{\text{sym}}$<sup>56</sup> |
| 1650 | $0.0214 \pm 0.0020$ | $0.0094 \pm 0.0020$ | (C=C) Amide I, Protein amide I absorption<sup>59</sup> |

Raman spectra of whole blood of the camel and human groups (Fig.4) show the difference in peak intensities at 540 cm^-1^ band, which is assigned to S-S signal. This band is reported as the sign of integral proteins of the cell membrane.^49,50^ The amount of the integral proteins was reported five times greater in camel erythrocytes compared to human erythrocytes.^18^ However, we observed that the spectral intensity of the 540 cm^−1^ band of the camels is twice as much of humans. The Raman signal at 714 cm^−1^ was assigned to phospholipid membranes. The current literature shows that RBCs membrane structures are different in human and camel species. Our data indicated that Raman intensity at 714 cm^−1^ is higher in camels than in humans. Such a difference may point out the structural difference between these two cell types. The two wavenumbers in the spectrum, 838 cm^−1^, and 945 cm^−1^, are absent for humans, while these are Raman-active for camels. The band 838 cm^−1^ was assigned to tyrosine^52,53^ and deformative vibrations of amine groups. The presence of the Raman signal at this peak for camels compared to humans remains inconclusive. The band, 945 cm^−1^, was assigned to polysaccharides and amylose. It was reported that camels have high blood glucose levels.^60^ In addition, the same study compared the amylase levels in camels as monogastric animals and humans as ruminant animals and found that intestinal amylase level is much higher in camels. Therefore, the main difference observed in 945 cm^−1^ for the two groups may stem from being a monogastric or ruminant animal.^60^ Intensities of the peaks at 1368 cm^−161^, which shows blood hemoglobin concentration level, are very close for the two groups.

## Conclusion

Cells are complex systems such that mechanical and chemical stimulations drive each other. With such complex and crowded lipid/protein compositions, it is challenging to attribute cells’ mechanical behaviors to specific lipid/protein types. Instead, deforming cells and observing their mechanical behaviors is another way of grasping cell mechanics. In this work, we aimed to investigate the most different cell types among the vertebrates, camel erythrocytes, compared to human erythrocytes. Examining the deformation characteristics of the cells using optical tweezers gave some results such that their deformability is one order of magnitude apart from each other. In addition, the whole blood analysis of the two species using Raman spectroscopy shed some light on the chemical content of these two types of cells, which are closely related to the mechanical features. However, we could not conduct single-cell Raman spectral measurements due to our technical limitations. Instead, we measured the whole blood. Nevertheless, the results we gathered from optical tweezers and Raman spectroscopy can fit together to reveal mechanochemical structural differences between these two species’ RBCs. Future efforts can explore the comparative nature of the human and camel RBCs by executing the single-cell Raman measurements as complementing cell deformation experiments at the single-cell level to have more satisfying results.

## Acknowledgements

This study was supported by a grant from the Directorate of Presidential Strategy and Budget of Turkey (Project Number: 2009K120520) and by a grant from Bogazici University Higher Education Council Scientific Research Projects (Project Number: 18143D).

## Author contributions statement

M.B.U. and U.P. conceived the experiment(s), T.P, M.H., and S.C. conducted the experiment(s), T.P, M.H., and S.C. analyzed the results, T.P. wrote the analysis codes, T.P. and M.H. created the figures. All authors reviewed the manuscript.

## Notes

### Competing Interest Statement

The authors have declared no competing interest.

## References

1. Anderson, H. L., Brodsky, I. E. & Mangalmurti, N. S. The evolving erythrocyte: red blood cells as modulators of innate immunity. The J. Immunol. 201, 1343–1351 (2018).

2. Ji, P., Murata-Hori, M. & Lodish, H. F. Formation of mammalian erythrocytes: chromatin condensation and enucleation. Trends cell biology 21, 409–415 (2011).

3. Snyder, G. K. & Sheafor, B. A. Red blood cells: centerpiece in the evolution of the vertebrate circulatory system. Am. zoologist 39, 189–198 (1999).

4. Ponder, E. The measurement of the diameter of erythrocytes. v.—the relation of the diameter to the thickness. Q. J. Exp. Physiol. Transl. Integration 20, 29–39 (1930).

5. Smith, J. E., Mohandas, N. & Shohet, S. B. Variability in erythrocyte deformability among various mammals. Am. J. Physiol. Circ. Physiol. 236, H725–H730 (1979).

6. Omorphos, S., Hawkey, C. M. & Rice-Evans, C. The elliptocyte: a study of the relationship between cell shape and membrane structure using the camelid erythrocyte as a model. Comp. biochemistry physiology. B, Comp. biochemistry 94, 789–795 (1989).

7. Perk, K. The camel’s erythrocyte. Nature 200, 272–273 (1963).

8. Perk, K., Frei, Y., Herz, A. et al. Osmotic fragility of red blood cells of young and mature domestic and laboratory animals. Am. journal veterinary research 25, 1241–1248 (1964).

9. Perk, K. Osmotic hemolysis of the camel’s erythrocytes. i. a microcinematographic study. J. Exp. Zool. 163, 241–246 (1966).

10. Fowler, M. Medicine and surgery of camelids (John Wiley & Sons, 2011).

11. Hoter, A., Rizk, S. & Naim, H. Y. Cellular and molecular adaptation of arabian camel to heat stress. Front. genetics 10, 588 (2019).

12. Yagil, R., Sod-Moriah, U. & Meyerstein, N. Dehydration and camel blood. ii. shape, size, and concentration of red blood cells. Am. J. Physiol. Content 226, 301–304 (1974).

13. Macfarlane, W., Morris, R., Howard, B. et al. Turn-over and distribution of water in desert camels, sheep, cattle and kangaroos. Nature 197, 270–271 (1963).

14. Warda, M., Zeisig, R. et al. Phospholipid-and fatty acid-composition in the erythrocyte membrane of the one-humped camel (camelus dromedarius) and its influence on vesicle properties prepared from these lipids. Deutsche Tieraerztliche Wochenschrift 107, 368–373 (2000).

15. Warda, M. et al. Proteomics of old world camelid (camelus dromedarius): Better understanding the interplay between homeostasis and desert environment. J. advanced research 5, 219–242 (2014).

16. Long, C. A. Evolution of function and form in camelid erythrocytes. Biophys Bioeng (2007).

17. Livne, A. & Kuiper, P. J. Unique properties of the camel erythrocyte membrane. Biochimica et Biophys. Acta (BBA)-Biomembranes 318, 41–49 (1973).

18. Eitan, A., Aloni, B. & Livne, A. Unique properties of the camel erythrocyte membrane: Ii. organization of membrane proteins. Biochimica et Biophys. Acta (BBA)-Biomembranes 426, 647–658 (1976).

19. Khodadad, J. K. & Weinstein, R. S. The band 3-rich membrane of llama erythrocytes: studies on cell shape and the organization of membrane proteins. The J. membrane biology 72, 161–171 (1983).

20. Ralston, G. B. Proteins of the camel erythrocyte membrane. Biochimica et Biophys. Acta (BBA)-Biomembranes 401, 83–94 (1975).

21. Condrea, E., Mammon, Z., Aloof, S. & De Vries, A. Susceptibility of erythrocytes of various animal species to the hemolytic and phospholipid splitting action of snake venom. Biochimica et Biophys. Acta (BBA)-Specialized Sect. on Lipids Relat. Subj. 84, 365–375 (1964).

22. Joshua, H. & Ishay, J. The haemolytic properties of the oriental hornet venom. Acta pharmacologica et toxicologica 33, 42–52 (1973).

23. Turner, J. C., Anderson, H. M. & Gandal, C. P. Comparative liberation of bound phosphatides from red cells of man, ox, and camel. Proc. Soc. for Exp. Biol. Medicine 99, 547–550 (1958).

24. Windberger, U., Auer, R., Seltenhammer, M., Mach, G. & Skidmore, J. A. Near-newtonian blood behavior–is it good to be a camel? Front. physiology 10, 906 (2019).

25. Ashkin, A. Acceleration and trapping of particles by radiation pressure. Phys. review letters 24, 156 (1970).

26. Ashkin, A., Dziedzic, J. M., Bjorkholm, J. E. & Chu, S. Observation of a single-beam gradient force optical trap for dielectric particles. Opt. letters 11, 288–290 (1986).

27. Ashkin, A., Dziedzic, J. M. & Yamane, T. Optical trapping and manipulation of single cells using infrared laser beams. Nature 330, 769–771 (1987).

28. Gieseler, J. et al. Optical tweezers—from calibration to applications: a tutorial. Adv. Opt. Photonics 13, 74–241 (2021).

29. McCauley, M. J. & Williams, M. C. Mechanisms of dna binding determined in optical tweezers experiments. Biopolym. Orig. Res. on Biomol. 85, 154–168 (2007).

30. Lincoln, B., Wottawah, F., Schinkinger, S., Ebert, S. & Guck, J. High-throughput rheological measurements with an optical stretcher. Methods cell biology 83, 397–423 (2007).

31. Mills, J., Qie, L., Dao, M., Lim, C. & Suresh, S. Nonlinear elastic and viscoelastic deformation of the human red blood cell with optical tweezers. Mol. & Cell. Biomech. 1, 169 (2004).

32. Guck, J. et al. The optical stretcher: a novel laser tool to micromanipulate cells. Biophys. journal 81, 767–784 (2001).

33. Agrawal, R. et al. Assessment of red blood cell deformability in type 2 diabetes mellitus and diabetic retinopathy by dual optical tweezers stretching technique. Sci. reports 6, 1–12 (2016).

34. Rancourt-Grenier, S. et al. Dynamic deformation of red blood cell in dual-trap optical tweezers. Opt. express 18, 10462–10472 (2010).

35. Bareil, P. B., Sheng, Y., Chen, Y.-Q. & Chiou, A. Calculation of spherical red blood cell deformation in a dual-beam optical stretcher. Opt. Express 15, 16029–16034 (2007).

36. Inanc, M. T. et al. Quantifying the influences of radiation therapy on deformability of human red blood cells by dual-beam optical tweezers. RSC Adv. 11, 15519–15527 (2021).

37. Jess, P. et al. Dual beam fibre trap for raman microspectroscopy of single cells. Opt. express 14, 5779–5791 (2006).

38. Sehgal, H., Aggarwal, T. & Salapaka, M. Characterization of dual beam optical tweezers system using a novel detection approach. In 2007 American Control Conference, 4234–4239 (IEEE, 2007).

39. Ott, D., Reihani, S. N. S. & Oddershede, L. B. Crosstalk elimination in the detection of dual-beam optical tweezers by spatial filtering. Rev. Sci. Instruments 85, 053108 (2014).

40. Raman, C. V. & Krishnan, K. S. A new type of secondary radiation. Nature 121, 501–502 (1928).

41. Rao, S., Bálint, Š., Cossins, B., Guallar, V. & Petrov, D. Raman study of mechanically induced oxygenation state transition of red blood cells using optical tweezers. Biophys. journal 96, 209–216 (2009).

42. Wood, B. R., Hammer, L. & McNaughton, D. Resonance raman spectroscopy provides evidence of heme ordering within the functional erythrocyte. Vib. spectroscopy 38, 71–78 (2005).

43. Raj, S., Wojdyla, M., Sanchez, M. M. & Petrov, D. Force and raman spectroscopy of single red blood cell. In Biophotonics: Photonic Solutions for Better Health Care III, vol. 8427, 842712 (International Society for Optics and Photonics, 2012).

44. Liu, R. et al. Novel single-cell functional analysis of red blood cells using laser tweezers raman spectroscopy: application for sickle cell disease. Exp. hematology 41, 656–661 (2013).

45. Deng, J., Wei, Q., Zhang, M., Wang, Y. & Li, Y. Study of the effect of alcohol on single human red blood cells using near-infrared laser tweezers raman spectroscopy. J. Raman Spectrosc. An Int. J. for Orig. Work. all Aspects Raman Spectrosc. Incl. High. Order Process. also Brillouin Rayleigh Scatt. 36, 257–261 (2005).

46. Feret, L. La grosseur des grains des matières pulvérulentes (Eidgen. Materialprüfungsanstalt ad Eidgen. Technischen Hochschule, 1930).

47. Pons, M.-N., Vivier, H., Delcour, V., Authelin, J.-R. & Paillères-Hubert, L. Morphological analysis of pharmaceutical powders. Powder technology 128, 276–286 (2002).

48. MATLAB. https://web.archive.org/web/20220729190040/ https://www.mathworks.com/help/images/ref/bwferet.html, (accessed: 29.07.2022).

49. Wesełucha-Birczyńska, A. et al. Human erythrocytes analyzed by generalized 2d raman correlation spectroscopy. J. Mol. Struct. 1069, 305–312 (2014).

50. Nakamoto, K. Infrared and Raman spectra of inorganic and coordination compounds, part B: applications in coordination, organometallic, and bioinorganic chemistry (John Wiley & Sons, 2009).

51. Stone, N., Kendall, C., Smith, J., Crow, P. & Barr, H. Raman spectroscopy for identification of epithelial cancers. Faraday discussions 126, 141–157 (2004).

52. Naseer, K., Amin, A., Saleem, M. & Qazi, J. Raman spectroscopy based differentiation of typhoid and dengue fever in infected human sera. Spectrochimica Acta Part A: Mol. Biomol. Spectrosc. 206, 197–201 (2019).

53. Siamwiza, M. N. et al. Interpretation of the doublet at 850 and 830 cm-1 in the raman spectra of tyrosyl residues in proteins and certain model compounds. Biochemistry 14, 4870–4876 (1975).

54. Naumann, D. Infrared and nir raman spectroscopy in medical microbiology. In Infrared spectroscopy: new tool in medicine, vol. 3257, 245–257 (SPIE, 1998).

55. Shetty, G., Kendall, C., Shepherd, N., Stone, N. & Barr, H. Raman spectroscopy: elucidation of biochemical changes in carcinogenesis of oesophagus. Br. journal cancer 94, 1460–1464 (2006).

56. Wood, B. R. & McNaughton, D. Raman excitation wavelength investigation of single red blood cells in vivo. J. Raman Spectrosc. 33, 517–523 (2002).

57. Tfaili, S. et al. Confocal raman microspectroscopy for skin characterization: a comparative study between human skin and pig skin. Analyst 137, 3673–3682 (2012).

58. Atkins, C. G., Buckley, K., Blades, M. W. & Turner, R. F. Raman spectroscopy of blood and blood components. Appl. spectroscopy 71, 767–793 (2017).

59. Malini, R. et al. Discrimination of normal, inflammatory, premalignant, and malignant oral tissue: a raman spectroscopy study. Biopolym. Orig. Res. on Biomol. 81, 179–193 (2006).

60. Osman, A. M. & El Khatim, M. M. S. Polysaccharidases of the camel (camelus dromedarius) intestine and pancreas. Comp. Biochem. Physiol. Part A: Physiol. 69, 429–436 (1981).

61. Virkler, K. & Lednev, I. K. Raman spectroscopic signature of blood and its potential application to forensic body fluid identification. Anal. bioanalytical chemistry 396, 525–534 (2010).

